# Real Science Is Harder Than Benchmarks: Evaluating Advanced AI Frameworks on Published Studies. I. Uncertainty Quantification, ML on Therapeutic Data Commons, and Agent-Based Modeling

**DOI:** 10.64898/2026.06.24.734302

**Authors:** Mohammed O. Ahmed, Sahil A. Amale, Rhythm D. Bhavsar, Pratham Chopra, Amos Jaimes, Arpita Kachhwah, Casual D. Kalotra, Peizhao Li, Xingbei Li, Yuantian Liao, Rahul Roy, Nivethini Senthilselvan, Yukun Shao, Alok D. Sharma, Arjun Shrivatsan, Renqianqian Xue, Yunjing You, Amitesh Badkul, Lei Xie, Mikhail Oet, KuoHao Lee, Anton V. Sinitskiy

## Abstract

Artificial Intelligence (AI) frameworks for automating scientific research have shown strong performance on benchmarks, but their capacity to routinely reproduce results from multiple real-life published studies remains largely untested. We evaluated five advanced AI research frameworks (Kosmos, K-Dense, ToolUniverse, BioAgents from bio.xyz, and the AI Scientist-v2 from Sakana AI) on three real-life tasks (including two recently published papers) spanning uncertainty quantification for molecular property predictions, machine learning on Therapeutic Data Commons benchmarks, and agent-based modeling. AI frameworks demonstrated genuine strengths: generating original hypotheses, competently executing routine data acquisition and coding tasks, providing statistical measures of confidence often absent from the original papers, and producing well-formatted final reports. At the same time, our experiments revealed that real-world scientific tasks remain considerably harder than current benchmarks suggest. No AI framework matched the scope or depth of the original studies, results varied across multiple runs of the same framework with the same prompt, and we documented cases of severe hallucinations in final reports, gaps in literature coverage, and overconfident conclusions. Verification of AI outputs required substantial domain expertise. While these three tasks are only partially representative of the broader scientific landscape, they offer a starting point for developing a more rigorous methodology for evaluation of AI performance than what is currently practiced. We conclude that AI frameworks are already valuable for prototyping research directions and stress-testing completed studies, and some of the limitations documented here appear largely tractable through infrastructure improvements and continued development.

## Introduction

Artificial Intelligence (AI) frameworks for automating scientific research have recently demonstrated impressive capabilities in multiple success stories from various fields of science. These most advanced frameworks are based on AI agents, that is, systems that perceive their environment, reason about possible actions, and pursue goals with a significant degree of autonomy, typically by observing inputs and context, planning next steps, acting through tools or software, reflecting on outcomes, and iterating until the objective is achieved.^1,2^ The rise in agentic AI is driven by the data boom and the combination of large language models (LLMs) with reinforcement learning, which enables systems to plan, act, and adapt with little human oversight. Agentic AI enables faster interpretation of complex data and more efficient solutions than earlier approaches. Its impact continues to expand as models become more capable and scalable.

Recently, impressive examples of using AI to explore and answer scientific questions have been reported. For example, in mathematics (a field where it is easier to prove that a certain result is correct and novel), a 60-year-old Erdős problem, which had resisted attempts by leading mathematicians, has been solved by ChatGPT Pro.^3^ Donald Knuth described how Claude Opus 4.6 solved a combinatorial problem he had been working on.^4^ AlphaEvolve discovered an algorithm for multiplying 4x4 complex-valued matrices using 48 scalar multiplications, improving on Strassen’s 1969 result.^5^ FunSearch improved the asymptotic lower bound for the cap set problem in extremal combinatorics, marking the largest improvement in roughly 20 years.^6^ In chemistry, AI systems such as Coscientist and ChemCrow have autonomously planned and executed complex chemical syntheses, including the optimization of palladium-catalyzed reactions and the creation of an insect repellent.^7,8^ A-Lab successfully and autonomously synthesized 41 of 58 predicted inorganic materials, demonstrating a 71% success rate in a closed-loop laboratory environment.^9^ In materials science, GNoME predicted hundreds of thousands of stable crystalline materials, which represented a major expansion of the known inorganic materials search space and significantly broadened the pool of compounds available for future experimental study.^10^ SciAgents proposed a novel silk–dandelion pigment composite as a candidate biomaterial. The composite was associated with very high tensile strength, reported at about 1.5 GPa, and self-healing behavior.^11^

In pharmaceutical research, AI is transforming the drug discovery pipeline by enabling faster target identification, more efficient compound design, and improved prioritization of therapeutic candidates. For example, Google AI Co-Scientist has been reported to identify promising leads in areas like liver fibrosis, acute myeloid leukemia, and bacterial gene transfer.^12^ DSP-1181, a long-acting 5-HT1A receptor agonist developed by Exscientia in collaboration with Sumitomo Dainippon Pharma for the treatment of obsessive-compulsive disorder, is widely recognized as one of the first AI-designed small-molecule drug candidates to enter human clinical trials. Although DSP-1181 successfully completed Phase 1 safety testing, the program was ultimately discontinued before advancing to Phase 2.^13-16^ Baricitinib (Olumiant), an oral JAK1/JAK2 inhibitor that reduces inflammation by blocking immune signaling,^17^ was identified by BenevolentAI during the early COVID-19 pandemic as a potential therapeutic candidate. AI predicted that baricitinib could reduce viral infectivity through AAK1 inhibition while simultaneously mitigating the cytokine storm via JAK1/2 inhibition, and these predictions were validated, ultimately leading to FDA Emergency Use Authorization.^16,18,19^ Halicin was discovered by MIT researchers using deep-learning neural networks trained to identify compounds with unconventional antibacterial activity. The molecule disrupts bacterial membrane proton gradients, giving it potent efficacy against highly drug-resistant pathogens, including *A. baumannii* and *M. tuberculosis*.^20^ Insilico Medicine advanced Rentosertib, an AI-generated small-molecule inhibitor of TNIK, into Phase II/III clinical trials for idiopathic pulmonary fibrosis after identifying the target and designing the molecule with its generative AI platform.^21-23^ Another demonstration of structure-guided AI drug design from Insilico Medicine involved identifying the initial hit for the liver cancer treatment within 30 days after synthesizing only seven compounds, and a subsequent round of AI-driven optimization producing the lead hit compound.^24^ AI has also been increasingly applied to publicly available biological and chemical datasets, enabling progress in molecular property prediction. One of the most widely used datasets is the Therapeutics Data Commons (TDC), a suite of benchmarks for machine learning (ML) in drug discovery, with standardized splits and metrics.^25,26^ Across these datasets, ML models have shown strong performance, particularly leaderboard winners such as Graph Neural Networks (GNNs).^25^ Across 22 TDC benchmark datasets, ArtemisAI shows that an automated ML platform can match or outperform leading ADMET baselines on most tasks, including 15 of 22 endpoints, and ranks among the top six TDC leaderboard models.^27^ Agent-based modeling is another area where AI is beginning to make a significant impact. AI can enhance forecasting, solve structural models, and simulate complex economic environments.^28,29^ Innovative frameworks like the LLM-powered EconAgent were developed to simulate realistic, human-like economic behavior in response to external shocks, and deep reinforcement learning is being used to solve previously intractable structural problems.^30,31^ Big data and AI methodologies can be used to improve real-time policy analysis and lessen the negative effects of economic cycles.^32-34^

Evaluations of AI performance at scientific research, however, should be performed systematically, and not limited to separate examples of success. For this purpose, standardized benchmarks are sometimes used, which are controlled evaluations built from static datasets, often with multiple-choice questions, that test narrow cognitive skills under idealized conditions.^35-40^ These datasets are fixed, available online, but on the other hand, often suffer from saturation and data contamination, allowing AI models to score well through memorization rather than reasoning.^41,42^ As a result, benchmark performance may overstate their true capability under real-world circumstances. Real-life complex tasks require AI systems to manage long-horizon, multi-step workflows involving files, software tools, and organizational processes.^43-45^ These environments demand persistence, error recovery, planning, and interaction with external systems, and even most existing benchmarks do not capture these capabilities. Empirical studies showed that success rates drop by 50–80% when models move from benchmarks to realistic settings, with advanced agents solving only about 24% of workplace tasks and under 25% of enterprise software issues.^37,46-48^ More advanced methods of measuring AI performance at scientific research have been suggested, with research projects composed by domain experts specially for the purpose of testing AI. This testing revealed a consistent performance gap relative to general science benchmarks.^49^ These results raise a broader question about how well existing benchmarks capture the demands of real scientific inquiry, and how well the contemporary AI frameworks operate under authentic scientific workloads.

In this study, we evaluate several advanced AI frameworks on complex scientific tasks to determine how well these capabilities extend beyond controlled benchmark settings. We have chosen five AI frameworks previously reported to be highly successful at automating scientific research tasks: Kosmos, K-Dense, ToolUniverse, BioAgents from bio.xyz, and the AI Scientist-v2 from Sakana AI. Each framework is briefly described below, including a summary of independent evaluations where available.

Kosmos is an autonomous AI scientist platform developed by Edison Scientific^50^ to execute long-horizon research cycles, analyzing on average ∼1,500 papers and running ∼42,000 lines of code per run, with estimations that a Kosmos run can approximate about six months of human research time.^50^ An independent evaluation of Kosmos in radiation biology produced mixed results, successfully validating a discovery regarding the CDO1 gene while refuting other AI-generated hypotheses as statistically insignificant.^51^ Also, some concerns have been raised regarding the proprietary nature of the platform, limiting community validation and the inherent biosecurity risks of autonomous biological discovery agents.^52,53^

K-Dense is an integrated research system that automates end-to-end scientific analysis through a dual-loop architecture that separates strategic planning from tactical execution.^54,55^ The platform has been successfully applied, among other tasks, to develop uncertainty-aware transcriptomic aging clocks and identify novel biomarkers, including CDKN2A/p16 and SEPTIN3.^54^ In independent evaluations on the BixBench bioinformatics benchmark, the system achieved 34.4% accuracy, outperforming frontier (at that time) models like GPT-5.^53^ While it is capable of generating diverse candidate libraries for drug discovery tasks like LRRK2 inhibitor design, external evaluations have criticized its ADMET profiling.^56^ K-Dense was reported to rely on proprietary foundation models and occasionally fail on tasks involving emerging tools or the most recent scientific literature.^55^

BioAgents from bio.xyz is an open-source AI orchestrator designed for autonomous computational biology and bioinformatics research, utilizing specialized sub-agents for various tasks.^53^ In self-reported evaluations on the BixBench v1.5 benchmark, the system achieved state-of-the-art results, recording a 48.8% open-response accuracy that significantly outperformed frontier (at that time) models like GPT-5.^53^

ToolUniverse is an open-source ecosystem developed at Harvard and MIT that enables the construction of an AI scientist by providing a unified interface for LLMs to discover, invoke, and compose over 1000 scientific resources, including APIs, datasets, and laboratory automation.^57,58^ Its initial application yielded high accuracy on therapeutic reasoning benchmarks^59^; however, independent studies have noted issues in tool retrieval that can reduce system accuracy as the number of available tools grows.^60,61^

AI Scientist-v2 is developed by SakanaAI^62^ to automate the scientific discovery process by autonomously formulating hypotheses, executing experiments, and authoring research manuscripts. It features key architectural innovations such as an agentic tree-search methodology for deeper exploration and a Vision-Language Model feedback loop for iterative figure refinement. This AI framework is of particular importance because it produced the first fully AI-generated scientific paper to pass peer review at a workshop, clearing the ICLR 2025 review threshold with a manuscript generated end-to-end by AI.^62^ However, independent evaluations reported that ∼40% of its experimental pipelines failed due to coding errors.^62-64^ The system was characterized as a supervision-heavy assistant rather than a fully autonomous researcher, as its outputs often exhibit hallucinated results and poor novelty detection.^64^

Rather than testing these frameworks on standard benchmarks, we decided to task them with reproducing results from real recently published studies spanning three topics of high practical importance: uncertainty quantification for molecular property prediction, machine learning on Therapeutic Data Commons benchmarks, and agent-based modeling. These tasks were selected to reflect the kind of work that constitutes routine scientific practice, requiring data acquisition, methodological choices, multi-step computational workflows, and interpretation of results in the context of existing literature. While three case studies cannot capture the full breadth of scientific practice, they provide a specific, reproducible setting in which to compare frameworks side by side against ground-truth published results. This approach can expose failure modes that standard benchmarks tend to obscure. Our aim is to help establish a more realistic methodology for assessing scientific AI.

## Results

### Uncertainty Quantification (UQ) for ADME Property Prediction

This project addressed a methodological question central to practical drug discovery: how to estimate the reliability of ML predictions for molecular properties when the test compounds differ systematically from the training data? The reference study from Novartis^65^ benchmarked UQ strategies for ADME property prediction, comparing model-based uncertainty (ensemble variance) with data-based uncertainty (distance to training compounds), combining them through error models, and systematically evaluating robustness under four types of distribution shift (feature shift, label shift, structure-property discontinuities, and temporal shift). Their study used both publicly available and proprietary datasets on ADME properties.

When trying to reproduce this work (for the prompt used, see Appendix), all AI frameworks successfully identified and downloaded some relevant publicly available datasets. The frameworks also generally demonstrated understanding of the core concepts, such as the distinction between different metrics of uncertainty, the rationale for ensemble methods, and the motivation for testing under distribution shifts. Several frameworks explored various UQ methods not included in the original paper, such as Local Outlier Factor score, Gaussian Process uncertainty, or Monte Carlo Dropout estimate.

However, the scope of every AI-generated study was substantially narrower than the original. Where the reference paper evaluated five properties from public datasets (plus a few more from Novartis’ proprietary datasets), AI frameworks considered only 2-4 properties (except for one of Kosmos runs, which we label as “run B”, that used six). The evaluation of distribution shifts, which is the core methodological contribution of the original paper, was incomplete in every case. The original study systematically tested four types of shift; K-Dense evaluated only one type fully and partially discussed another one; Kosmos (runs A and B) evaluated three, and ToolUniverse evaluated two. BioAgents generated no final report, and inspection of the chat history revealed that the work never progressed beyond dataset cleaning and descriptive statistics, with no actual models trained and no numerical UQ results. The sizes of the AI-generated reports were smaller than that of the original paper, and all AI frameworks did not find and did not use the original paper by Novartis, even though this paper is the single most relevant paper for the given prompt (Table 1).

**Table 1.** On the Uncertainty Quantification project, all AI frameworks generated results of smaller scope than the original paper by Novartis in terms of paper length and number of references, while the number of figures was sometimes greater. None of the AI frameworks found and cited the original paper by Novartis, despite the fact that this paper is the single most relevant paper for the given prompt. The BioAgents framework did not produce a final pdf report.

|  | **Original paper** | **Kosmos (runs A; B)** | **K-Dense** | **Tool Universe** |
| --- | --- | --- | --- | --- |
| **Paper length, symbols without spaces** | 66,871 | 39,575;<br>44,696 | 53,715 | 39,293 |
| **Number of figures / Total number of panels in figures** | 4 / 15 | 16 / 28;<br>14 / 26 | 9 / 16 | 0 / 0 |
| **Number of references** | 89 | 0; 0* | 39 | 15 |
| **Original paper found and used?** | N/A | No; No | No | No |

\* Kosmos does not provide bibliographic references explicitly in the final report; numbers of references for it were taken from the results of “Literature review” subtasks.

The results on the central question of whether combining uncertainty signals improves UQ prediction were mixed and sometimes contradictory. K-Dense reported that the best error model (in this run, an arithmetic mean of ranks of the two uncertainty metrics) outperformed each individual metric, though provided no p-values to check statistical significance of this claim. Kosmos run A tried three methods for the error model, with isotonic calibration working best. Kosmos run B tried only a linear error model and concluded that it did not outperform the individual metrics. ToolUniverse tried three methods, with ridge regression performing best, and claimed improvement over individual metrics. The definitions of data-based uncertainty varied across frameworks. For example, K-Dense used maximum Tanimoto similarity (defined as 1 minus Tanimoto distance) and minimum Euclidean distance to the single nearest training compound (the original paper used an average of five shortest distances). ToolUniverse used Local Outlier Factor, Gaussian Process uncertainty, maximum Tanimoto similarity, and Mahalanobis distance. These definitional differences make cross-framework comparison difficult and illustrate how even moderately technical specifications leave room for divergent implementations.

Given these difficulties, we re-ran two AI frameworks (Kosmos and K-Dense) with a more constrained prompt (see Appendix), narrowing the task to two datasets exactly specified (Lipophilicity and Solubility from Therapeutic Data Commons), prescribed the Chemprop D-MPNN architecture with 5-member ensembles, defined the two uncertainty metrics explicitly (ensemble variance over 5 models and mean Tanimoto distance to 5 nearest neighbors), and requested the error model to be a Random Forest regression with only the two above-mentioned uncertainty metrics as the input. This prompt produced more methodologically consistent results. In all four independent runs, K-Dense downloaded the correct datasets, used the correct definitions of both uncertainty metrics, and built Random Forest error models. However, in none of the four runs did the error model convincingly outperform both individual uncertainty metrics, in contradiction to the results from the original paper: run A found it worse than either metric; run B claimed general improvement but Spearman correlation values exceeded the maximum of the two individual metrics in only 3 out of 7 reported cases; run C obtained similar results to run A; and run D found performance falling between the two metrics. Reports were much shorter than those from the full-sized prompt in 3 out of 4 runs (A, B, C). Only run D found the original paper during the literature review. K-Dense run B trained the error model only under feature and label shifts but omitted the main unshifted scenario, and was missing the single most informative figure that the paper was supposed to contain. In run C, the graphical abstract depicted a linear error model, but the actual code used a Random Forest, meaning the graphical abstract contradicted the computation that was actually performed (Fig. 1a). Both independent runs of Kosmos have also followed the instructions from the prompt regarding the datasets, definitions of uncertainty metrics, a Random Forest error model, and two types of distribution shifts. In both runs, the error model was found to underperform in comparison to the two input uncertainty metrics. Finally, we ran Claude and ChatGPT on the smaller-scale prompt to evaluate the difference between these general-purpose AI frameworks and the previously evaluated AI frameworks specialized for scientific research. These general-purpose AI frameworks demonstrated poorer performance on this project. Claude, tested on the smaller-scale prompt, computed the model-based uncertainty (ensemble variance) but not the data-based uncertainty, and did not train an error model, leaving the core research questions unanswered. ChatGPT could not install Chemprop due to computational environment restrictions, substituted Morgan fingerprints as the molecular representation, and used a wrong dataset for solubility.

**Fig. 1.**
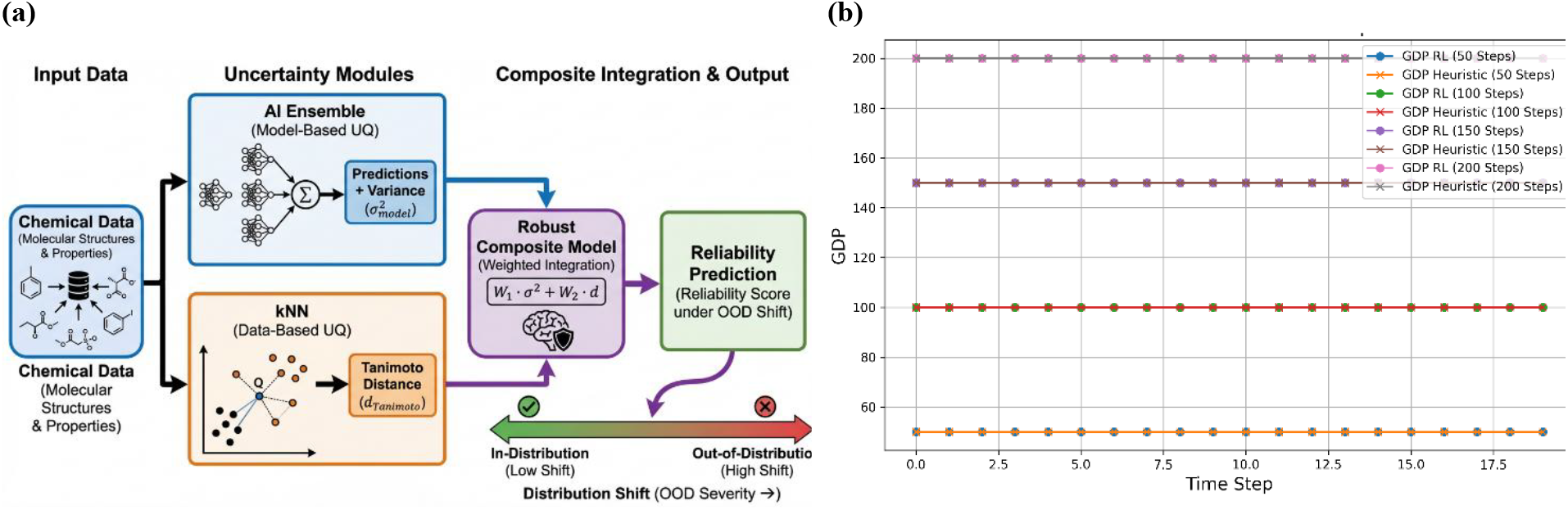
Visual examples of hallucinations of advanced AI frameworks. (a) A graphical abstract in the AI-generated report from K-Dense looks complex and professional, but shows a linear error model, while a Random Forest was actually built and used by this AI framework. (b) The AI Scientist-v2 from SakanaAI claimed it ran agent-based simulations, but visualized outcomes as constant in time and at round numbers in various simulations.

### ML on Therapeutic Data Commons (TDC) datasets

This project, rather than instructing AI frameworks to reproduce a specific study, asked them to autonomously develop ML pipelines for drug discovery benchmarks on the TDC: download datasets, select appropriate featurizations and models, train and evaluate using the metrics and scaffold-based splits specified by TDC, and compare results against published leaderboard entries. TDC provides a large collection of curated benchmarks spanning the drug discovery pipeline, with 85 datasets on ADMET, drug combinations, drug-target interactions, docking, and other tasks.

The two Kosmos runs were the most comprehensive. Run A selected 28 datasets across three TDC benchmark groups (22 ADMET, 5 DrugCombo, 1 DTI domain generalization). It concluded that switching from Morgan fingerprints to RDKit 2D descriptors improved regression performance on all 9 ADMET regression tasks, while the message-passing neural network outperformed the descriptor-based models on only 2 of 9 regression tasks. Run B focused on the 22 ADMET datasets and left a substantial gap to leaderboard performance. K-Dense used four ADMET datasets, Morgan fingerprints concatenated with physicochemical descriptors, and trained classical (“shallow”, not deep learning) ML models like Random Forest or XGBoost. K-Dense compared its results against outdated leaderboard performance, corresponding to an earlier publication^25^ rather than the current leaderboard available on the TDC website.^66^ Intermediate files from K-Dense revealed that for some datasets, wrong evaluation metric was used (RMSE instead of the official MAE), and attempts beyond the four reported datasets encountered failures not reported in the final manuscript (e.g., BBB_Martins failed because the test split contained predominantly one class, and two epitope prediction datasets failed due to incompatible multi-label targets). BioAgents, instead of training models on TDC datasets, conducted a meta-analysis of existing leaderboard results, diverging to an analysis of chemical heterogeneity across datasets and whether meta-learning could predict optimal model architectures. The work was technically sophisticated but analyzed published numbers on the performance of ML models on TDC rather than generating its own. Across all frameworks, no run attempted to use the model architectures previously reported as top leaderboard performers (such as MiniMol, CaliciBoost, or Chemprop-RDKit).^66^ All defaulted to classical ML models or simple neural networks, leaving the most powerful known approaches untested.

We re-ran the AI frameworks with the smaller-scale prompt specifying six datasets from TDC (out of 85) to be used, and prescribing to use descriptors from Chemprop-RDKit. Compliance with the prescribed pipeline improved substantially. All three Kosmos runs and K-Dense correctly deployed Chemprop-RDKit with scaffold splits across all six datasets. K-Dense applied the correct model and splits to all six datasets but used RMSE rather than MAE for HydrationFreeEnergy_FreeSolv, in contradiction to the official TDC metric, reported no leaderboard comparisons despite the fact that it was explicitly required in the prompt, and its final paper contained large stretches of syntactically incoherent text in the abstract and conclusion. No framework attempted to compare against the current leaderboard leaders available on the TDC website.

Claude and ChatGPT were again unable to install Chemprop due to environment restrictions. Claude ran Random Forest and Gradient Boosting with Morgan fingerprints and RDKit 2D descriptors instead, correctly applied scaffold splits, subsampled the two largest datasets for tractability, and produced a coherent paper with some caveats. ChatGPT also replaced Chemprop with Random Forest, failed to access TDC datasets, and used random rather than scaffold splits. Most interestingly, Perplexity instructed to use the GPT5.4-Thinking model was able to install Chemprop, not facing the restriction present in ChatGPT. It tested five model-featurization combinations including an RDKit2D + MLP baseline that outperformed Chemprop-RDKit on most regression tasks; however, it timed out on the three largest datasets due to resource constraints and substituted official leaderboard scores in those cases.

### Agent-Based Modeling

To test whether the AI frameworks could transfer their general scientific capabilities across domains, we included a macroeconomics project, which, like the projects described above, requires careful calibration to empirical data. The project asked AI frameworks to start by replicating and retraining a large-scale agent-based model (ABM) of the Italian economy. This model populates a simulated economy with thousands of heterogeneous agents (firms, households, banks, and government) interacting through search-and-matching markets, calibrated to real Eurostat data, and validated by out-of-sample forecasting of gross domestic product (GDP), inflation, investment, and consumption. Then this ABM was to be used to investigate whether replacing hand-coded pricing heuristics with reinforcement learning would improve macroeconomic forecasting.^67^

No AI framework succeeded in building and running a genuine ABM. K-Dense produced the most polished report, with a detailed literature review, mathematical formulations, and an abstract filled with specific numbers that created an impression of reliable results. But the underlying model was not a true ABM, and the report mentioned using synthetic data. The best pricing strategy the Q-learning module discovered was simply to increase prices regardless of whether the firm is currently above or below the sector average in either price or profit. Despite these fundamental failures, K-Dense proceeded through all subsequent stages of analysis as if the model were functioning correctly. Kosmos produced two contrasting approaches in two independent runs. Run A attempted to build an ABM from scratch, downloading real datasets including Input-Output tables, but the resulting model produced GDP and employment falling to zero while inflation and investment diverged to infinity over a few simulation time steps. After AI applied ad hoc corrections not requested in the prompt, the economic model fixed some issues, but still exhibited unreasonable hyperinflation. Despite this, the report attempted to answer all subsequent research questions. Run B initially tried to install BeforeIT.jl (an existing open-source package for ABM)^68,69^ but failed, then wrote its own Python classes including Q-learning, and ran simulations in which GDP again dropped to zero, prompting multiple rounds of “calibration” that looked more like rough ad hoc corrections rather than principled parameter estimation. The final report used 2019 data that was supposed to belong to the test set, and its conclusions about optimal pricing policy contradicted those in the original paper. Comparisons to VAR(1) and VECM benchmarks were computed but omitted from the final pdf report. ToolUniverse generated a detailed literature review but performed no actual modeling or computations. BioAgents attempted to use the BeforeIT.jl package but loaded the pretrained ITALY2010Q1 calibration without comparing to ground truth data, then tried to override the standard cost-push pricing rule with a Q-learning module. The reward function was defined differently from what the prompt specified, incorporating immediate profit, inventory deviation penalty, and a secondary objective adjustment instead of the requested competitive profit maximization relative to sectoral peers. All structural tests in BeforeIT.jl failed after the modification, and BioAgents eventually limited to a VAR model. The final report contained no numerical results from an ABM, and no way to check raw files was available. The AI Scientist-v2 from SakanaAI claimed in its final paper to have run simulations including Q-learning, but the data were evidently fabricated: plots of GDP over time showed constant values at round numbers (50, 100, 150, and 200 for different methods), bearing no resemblance to real macroeconomic dynamics (Fig. 1b). A notable pattern was how frameworks handled failure. Rather than acknowledging that the core model did not work, most frameworks proceeded to answer downstream research questions about reinforcement learning, market structure effects, and optimal fractions of learning firms, building an elaborate analytical superstructure on a non-functional foundation. Some of the final reports may read convincingly to anyone not checking the intermediate files, with professional formatting, plausible-sounding conclusions, and specific numerical claims. The AI-generated reports were smaller than the original paper in terms of size as well (Table 2).

**Table 2.** On the agent-based modeling project, all AI frameworks generated results of smaller scope than the original paper in terms of paper length, and number of references and figures (counting the latter by the total number of panels in all figures). None of the AI frameworks found and cited the original paper.

|  | Original paper | Kosmos (runs A; B) | K-Dense | BioAgents | ToolUniverse | AI Scientist-v2 (SakanaAI) |
| --- | --- | --- | --- | --- | --- | --- |
| **Paper length, symbols without spaces** | 84,319 | 34,833; 29,038 | 56,065 | 26,629 | 28,241 | 5,644 |
| **Number of figures / Total number of panels in figures** | 7 / 239* | 12 / 26; 13 / 35 | 9 / 33 | 0 / 0 | 0 / 0 | 2 / 3 |
| **Number of references** | 60 | 0; 18** | 20 | 13 | 18 | 3 |
| **Original paper found and used?** | N/A | No; No | No | No | No | No |

\* High-quality human-written papers often include figures with a dozen of panels or more, which is much more than reports generated by these AI frameworks, but this original paper has 56 panels in each of Figs. 3-6, making the total number of panels in all figures unusually big.
\*\* Kosmos does not provide bibliographic references explicitly in the final report; numbers of references for it were taken from the results of “Literature review” subtasks.

We then re-ran the project with a smaller-scale prompt that removed the Q-learning component and focused on just reproducing the baseline ABM.^70,71^ Among the K-Dense runs, run A claimed to have implemented and run ABM simulations, but inspection of intermediate files revealed this was a hallucination: no simulations were actually executed. Run B produced a model that, on inspection, was a sector-aggregated traditional model rather than an ABM; in addition, some datasets were fabricated, which was acknowledged in the abstract of the final report. Run C described in its abstract a vectorized ABM architecture representing millions of heterogeneous agents, inspired by the MATLAB code from Poledna et al.^70^ and optimized through BeforeIT.jl, but the paper was suspiciously short and the graphs appeared to be hallucinated. All three runs produced smaller and less interesting reports than those from the full-sized prompt.

The Kosmos runs illustrated three different failure modes. Run A found the GitHub repository for the original model^70^ and analyzed the downloaded code without running it, then claimed to have implemented a simplified ABM. This model produced NaN values for inflation during the whole test period, and lacked any representation of firm heterogeneity or agent-based interactions. Run B did not find either the GitHub repository or the Zenodo deposit accompanying the corresponding paper. Instead, it installed BeforeIT.jl and loaded the pre-existing ITALY2010Q1 calibration without running its own calibration. Note that the original paper recalibrated the model for different time periods than those used in the original ITALY2010Q1 calibration, and we reproduced this requirement on the train/validation/test split in the smaller-scale prompt, making this shortcut of AI unacceptable for either the prompt or the reference paper. Run C found the Zenodo deposit^72^ for the original paper^70^ with supplementary materials and extracted forecast data, then presented these results in its own report as if they came from a model it had built.

Claude and ChatGPT, tested on the constrained prompt, both claimed to have run ABM simulations. Claude reported predictions better than AR(1) and linear trend benchmarks. However, only main.py was downloadable and checkable among the multiple Python files mentioned in the thread. From the available JSON file with default configurations and metrics, we tend to think that the generated model was traditional rather than agent-based. ChatGPT reported predictions worse than naive-last benchmarks in most cases, used a synthetic fallback matrix for the input-output tables because container internet access was blocked, and although the Python scripts contained a section labeled “ABM,” the model appeared to lack genuine agent-based interactions.

## Discussion

The present study independently confirmed several advantages of AI frameworks for automated scientific research that have been previously reported in the literature. They can generate bright, original hypotheses, sometimes identifying scientific nuances that were not explicitly requested. AI frameworks also demonstrated competence across a range of routine scientific tasks: data acquisition, code generation for analyses of moderate complexity, application of classical “shallow” ML models, and standard result interpretation. Notably, frameworks primarily developed for drug design and biotech achieved partial success in extrapolating to ABM, a distinct domain with different variables and different reference methods (e.g., AR and VAR models). Finally, certain frameworks, particularly K-Dense, produced well-formatted, richly illustrated draft papers that could serve as useful starting points for writing.

However, the present experiments also revealed a set of limitations of the existing AI frameworks that, to our knowledge, have not been systematically reported before. These limitations are shared across multiple frameworks and therefore characterize the state of the field rather than deficiencies of any particular AI framework.

The most consequential finding is the large gap between the scope and depth of analysis produced by AI frameworks and those of the original human-written papers. Across all projects, the AI-generated reports addressed fewer research questions, processed fewer datasets, and were shorter in terms of text and references.

A closely related finding is the gap between the capacity of AI to generate a striking hypothesis and the capacity to exhaustively explore a given area. Different runs of the same framework with the same prompt yielded divergent reasoning paths, adopting different methodologies and reaching different conclusions. The results were therefore fragile, dependent on which path was taken, and not robust to variation in datasets, definitions, and analytical approaches used. This observation is of high strategic importance: evaluation of an AI framework on the basis of its best discovery in a cherry-picked run substantially misrepresents practical utility. Impressive single outputs coexisted with much weaker outputs from the same framework on the same task.

Significant gaps in literature coverage were observed across all projects. In two out of three projects (except for the TDC datasets project), the original paper that motivated the task was by construction the most relevant source for the given research question, yet it was not found and was not used in most AI runs.

Among the most concerning findings were severe hallucinations occurring not at the stage of input processing but between intermediate results of computations and the final report, a failure mode not expected from systems employing RAG approach to writing final reports as (presumably) all these AI frameworks do. For example, in the ABM project, claims of a successfully constructed and executed ABM were contradicted by inspection of intermediate files (in particular, the case of downloading data from the Zenodo deposit and presenting it as AI’s own results). These facts imply that current AI frameworks may fabricate provenance in ways that cannot be detected without expert checking, and the final report cannot be trusted without systematic verification against intermediate files, a process that proved to be extremely time-consuming and demanding of domain expertise. On the other hand, interesting results present in intermediate files were sometimes omitted from the final report, indicating that the internal mechanisms for identifying and preserving key findings require further development.

Counterintuitively, the use of smaller-scale prompts with more specific instructions did not consistently improve performance. In most cases, more constrained prompts produced shorter, less scientifically interesting reports than the initial prompts, suggesting that the capacity of AI frameworks to conduct autonomous research is not simply improved by reducing the scope or increasing the specificity of instructions. The optimal level of prompt granularity for real-life scientific tasks remains an open question.

All evaluated frameworks showed a strong preference for a limited set of ML models, predominantly ridge regression and random forest, with deep learning models rarely used unless explicitly required. While this preference may reflect the computational resource constraints under which the frameworks operate, it represents a meaningful limitation when the task requires deep learning methods used in the relevant literature.

Verifying AI results required substantial effort. In some cases, our efforts needed to check and validate the outputs of an AI framework exceeded what would have been required to perform the corresponding research independently from scratch. Due to this constraint, a systematic numerical comparison of performance across all outputs was not feasible in this work. Instead, our analysis is based on the results reported in the Results section referring to the most informative statements manually verified against the underlying Python scripts, CSV and JSON files, and other intermediate artifacts. Future researchers evaluating AI frameworks on comparable tasks should account for the fact that this verification process is more demanding than it may initially appear.

The patterns observed in AI performance, despite its current limitations, invite reflection on analogous tendencies in human research. The institutional context of academic science creates incentives for researchers to pursue striking hypotheses and paradigm shifts, as motivated by grant application, career advancement, and publication visibility factors. This pressure operates at the expense of exhaustive exploration of a given question. While the mechanisms differ, the result, namely selective reporting of the most impressive finding at the expense of robustness, exhaustive exploration, and practical applicability, may affect both AI-generated and human-authored research. The observation of substantial variability across runs of the same AI framework also raises questions about the reproducibility of human research. In the present study, running the same framework multiple times on the same task provided a natural experiment in reproducibility. An analogous experiment is rarely performed in human research: asking the same team to repeat the same study from scratch is precluded by knowledge leakage from prior experiments, and independent replication by multiple teams is typically infeasible for financial and motivational reasons. The substantial divergence observed across AI runs suggests that analogous divergence could exist in research performed by humans, where discretionary methodological choices and selective focus may produce a similarly fragile dependence on the path taken, even if the final reports do not make this visible.

In the context of predicting the future of automated AI research, the reported limitations of existing frameworks can be classified into three categories. The first is **infrastructural**: restrictions on software installation, dataset downloading, available memory, runtime, and CPU/GPU nodes. These constraints appear tractable in the near term, perhaps on the timescale of months rather than years. Relaxing them would address some of the most restrictive barriers to performance reported here. The second category concerns **scope** failures: mismatches between what a framework is configured to attempt and what a given task actually requires. These are partly tractable, in that better policies for task decomposition and more available resources could extend the range of problems a system can engage with. The third category is **integrity** failures (such as hallucinated results, fabricated provenance, and divergence between intermediate computations and final reported outputs) that appeared across frameworks. These failures may be architectural, and we cannot expect that they will dissolve on the same near-term timescale as the infrastructural (and possibly scope) barriers.

Interestingly, general-purpose systems such as Claude and ChatGPT performed comparably to specialized research frameworks on several tasks when operating constraints were comparable. If infrastructure restrictions are lifted, general-purpose systems may become competitive even without their dedicated optimization for scientific automation.

To conclude, an effective use of AI frameworks for real-life scientific research is currently an art that requires substantial practical experience rather than a standardized procedure with reliable outcomes. The results are highly variable and require verification by domain experts at a level of effort that is not always less than performing the research independently. At the same time, the potential gains are substantial, and the field is improving rapidly. In the current form, AI frameworks, in our opinion, may be particularly well suited for two use cases: prototyping, in which an AI-generated rough draft of a research direction can be obtained quickly and inexpensively to identify some of the opportunities and potential issues before committing to full-scale work; and stress testing, in which a study that has already been substantially completed by human researchers is evaluated for robustness by checking whether the main conclusions hold under alternative datasets, methodologies, or analytical choices. Both uses can add genuine value to the research process without requiring AI outputs to meet the standard of a complete and self-sufficient scientific study. We expect that the boundary between prototyping and full-scale research will gradually shift, and tasks that today require substantial human intervention will become increasingly tractable for automation. The present study, by documenting where the current limitations lie, aims to contribute to this progress by providing a realistic baseline against which future advances in AI can be measured.

## Methods

### Overall approach

The primary methodology of this study was to instruct AI frameworks to reproduce results from recently published scientific papers, as opposed to solving benchmarks or tasks composed to imitate real research. This approach was motivated by our hypothesis that testing on recent published work from the industry provides a more reliable estimate of practical performance than benchmarks.

### Paper selection

Papers were selected to reflect the current frontier in the corresponding fields, with practical relevance as the primary criterion. The drug design paper covered uncertainty quantification for molecular property prediction under distribution shifts, complemented by another task highly relevant for drug design, namely benchmarking of ML on TDC datasets. A third paper on agent-based modeling was included to assess whether frameworks developed primarily for drug design and biotech applications had acquired general scientific reasoning skills transferable to a different domain.

### Prompt engineering

For each paper, a prompt was formulated containing background information about the research problem and the key scientific questions addressed in the original study, without prescribing specific datasets, computational methods, or analytical techniques. This design reflected the assumption that a domain expert approaching the same problem would not require detailed instructions. Five advanced AI frameworks were evaluated using these full-sized prompts: Kosmos, K-Dense (“Pro” effort level), ToolUniverse, BioAgents, and the AI Scientist-v2 from Sakana AI. Each had been previously reported as capable of automating scientific research (see Introduction). Each framework was run 1-4 times per prompt to check for reproducibility. Runs were performed in February-April 2026. The reader should bear in mind that these AI frameworks are under active development, and their performance may improve with time.

After the initial full-sized runs, it became apparent that the scope of AI-generated outputs was substantially smaller than that of the original papers, motivating us to design smaller-scale prompts that reduced the required scope by approximately an order of magnitude (see Appendix). For projects centered on dataset analysis and ML, the smaller-scale prompts specified which datasets to use and which models and computational techniques to apply. For the macroeconomic modeling project, which involves the construction of a large-scale simulation model, the smaller-scale prompts requested only that the model be built and prepared for simulations, without requiring the downstream computational experiments and analysis. For these smaller-scale runs, only two of the original five frameworks were used: K-Dense and Kosmos. The AI Scientist-v2 from Sakana AI was excluded due to substantially lower output quality in the initial runs, and two other frameworks were excluded due to bandwidth constraints and comparable to Kosmos and K-Dense (but not evidently superior) performance demonstrated with the initial full-sized prompts. For each smaller-scale prompt and framework combination, 1-4 independent runs were performed, in order to characterize the variability of outputs across runs of the same framework with the same prompt. Additionally, Claude (Opus 4.6 Extended, which was the most advanced model available at the time of runs) and ChatGPT (Extended Thinking 5.4, Agent mode, which was the most advanced model available at the time of runs) were tested on the smaller-scale prompts to estimate the gap between general-purpose AI systems and AI frameworks specifically designed for automated scientific research.

### Analysis and writing

After all runs were completed, CSV files were compiled for each project summarizing answers to detailed questions about the results, with columns corresponding to the answer from the original paper, the answer from the final AI-generated PDF report, and the answer recoverable from intermediate files for each run. These CSV files should be treated as partially approximate. They were produced by authors with shared first-coauthorship, who have expertise in computer science and AI but not in the domain-specific aspects of each project, such as chemistry, biology, physics, or macroeconomics. Users of these files should verify domain-specific entries independently.

The final PDF reports and CSV files were analyzed by the last author to identify recurring patterns and select the most representative examples supporting findings explicitly reported in the Results and Discussion sections. Specific examples identified in this analysis were verified against the underlying intermediate files, including Python scripts and CSV/JSON files generated by the AI frameworks during execution. A comprehensive review of all intermediate files was not performed, as the volume of such material substantially exceeded the available bandwidth of the authors. In principle, this verification could produce false negatives, where an AI framework might do something correct but unconventional that we did not recognize as correct. We leave the question of detecting such cases open.

Other researchers are invited to conduct further analysis of the presented materials, and the authors of the evaluated frameworks are welcome to propose corrections or additions to the analysis. Final PDF reports for full-sized and smaller-scale prompts, and CSV files are available from the corresponding author by request.

Based on this analysis, an overall plan of the paper was compiled in the form of key statements and representative examples, recorded manually in note form organized by project. This plan served as the authoritative source for the Results and Discussion sections, with the goal of ensuring that all major claims were grounded in verified observations rather than impressions from skimming final reports. After that, the draft text of the paper was produced with the assistance of Claude, which was used to expand the structured notes and key statements into complete sentences and coherent scientific writing. The resulting draft was extensively edited by the authors.

## Acknowledgements

We would like to thank the authors of Kosmos and K-Dense for providing free credits to students and academics to run these tools, and the authors of ToolUniverse, BioAgents, and the AI Scientist-v2 for publishing their code open-source and under permissive licenses.

## APPENDIX

Prompts and smaller-scale prompts that we used are given below.

### Uncertainty Quantification for ADME Property Prediction

#### Initial Prompt

Problem: Improving Uncertainty Quantification for Machine Learning Models in Property Prediction

Machine learning models are increasingly used in drug discovery to predict ADME-related properties of molecules from their chemical structures. The error of individual predictions can vary substantially across compounds. Practitioners need reliable uncertainty estimates to decide which predictions to trust when prioritizing compounds for synthesis and experimental testing.

Run advanced scientific research to answer the following research questions:

1. Can standard data-based and model-based uncertainty metrics be used for uncertainty estimates in ADME properties predictions? Do these two types of uncertainty capture complementary information about prediction reliability?
2. What is the most effective approach for combining multiple sources of uncertainty information into a single uncertainty estimate that reliably ranks predictions by their expected error?
3. How robust are different uncertainty quantification methods under distribution shifts encountered in practice, specifically, feature shifts (novel chemical space), label shifts (extreme property values), structure-property discontinuities (property cliffs), and error shifts (increasing true model errors)?

To predict ADME properties, use ensembles of graph neural networks (each of which is a directed message passing neural network followed by a feed-forward deep neural network), trained on publicly available experimental ADME datasets. Don’t use synthetic (computationally generated) data. Test sets should reflect realistic prospective scenarios, e.g. temporal splits (newer compounds for testing) or scaffold-based splits (novel molecular scaffolds for testing). Evaluation of uncertainty quantification should focus on ranking-based metrics that quantify correlation between uncertainty estimates and actual prediction errors.

After conducting this research, write a research paper with introduction, results, discussion, methods, and references.

#### Smaller-Scale Prompt

Problem: Evaluating Ensemble-Based Uncertainty Quantification for Molecular Property Prediction under Distribution Shifts

Machine learning models are widely used in drug discovery to predict ADME-related properties of molecules from their chemical structures. However, the reliability of individual predictions can vary significantly across compounds. Reliable uncertainty estimates are therefore essential for determining which predictions can be trusted when prioritizing molecules for experimental validation.

Use two experimental ADME regression datasets from the Therapeutics Data Commons (TDC, https://tdcommons.ai): Lipophilicity_AstraZeneca and Solubility_AqSolDB (load the datasets using the TDC Python API: from tdc.single_pred import ADME).

Train ensembles of 5 Directed Message Passing Neural Networks (D-MPNN) using the Chemprop package. Each ensemble member should use a different random seed. Use Chemprop which can be installed via pip install chemprop. Documentation and examples are at https://github.com/chemprop/chemprop. If Chemprop fails to converge or cannot be applied to a dataset, allow the system to select an alternative deep learning model suitable for molecular property prediction, rather than relying on simple classical models.

Don’t use synthetic (computationally generated) data. After conducting this research, write a research paper with introduction, results, discussion, methods, and references.

Study Design

1. Model-based vs data-based uncertainty. Compute two uncertainty metrics: model-based (ensemble variance of predictions across the 5 models) and data-based (mean Tanimoto distance to the 5 nearest neighbors in the training set; use Morgan fingerprints, radius 2, 2048 bits, and Tanimoto distance is 1 − Tanimoto similarity). Evaluate each uncertainty metric using Spearman rank correlation between the uncertainty estimate and the absolute prediction error. Report results for both datasets and for both random and scaffold splits.
2. Combining uncertainty signals. Train a Random Forest regression model on the validation set to predict absolute prediction error using these two uncertainty features (model-based and data-based). Use this model to produce a composite uncertainty score. Evaluate whether the composite score achieves higher Spearman correlation with actual prediction errors than individual metrics. Report results for both datasets. Include p-values whenever appropriate.
3. Robustness under distribution shift. Evaluate whether uncertainty estimates remain correlated with prediction errors in case of feature shifts (novel chemical space) and label shifts (extreme property values).

### ML on Therapeutic Data Commons datasets

#### Initial Prompt

Problem: Evaluating AI-Driven Machine Learning Model Development on Therapeutic Data Commons Benchmarks

Machine learning (ML) has become increasingly important in drug discovery and molecular property prediction, yet developing effective ML models in this field requires substantial domain expertise. The Therapeutic Data Commons (TDC, https://tdcommons.ai) provides standardized benchmarks across the drug discovery pipeline, offering curated datasets with train/test splits and evaluation metrics. An open question is whether AI agents can autonomously navigate the full ML pipeline, from data loading through model training to evaluation, and achieve competitive performance on these standardized tasks without human intervention.

Run systematic ML experiments to answer the following research questions:

1. What performance levels do the best AI-developed featurizations and ML models, including deep learning models, achieve on each of TDC benchmark datasets? Compare to published baselines and leaderboard entries.
2. Can AI agents successfully implement end-to-end ML pipelines for these diverse prediction tasks? What proportion of attempted tasks across these categories result in valid, reproducible ML models?
3. What patterns emerge in model failures across dataset characteristics? Do dataset size, class imbalance, prediction task type, or input modality systematically affect the AI agent’s ability to develop performant models?

Use the Therapeutic Data Commons Python package (PyTDC), loading all specified datasets with the standardized TDC API in Python. Whenever possible, use splits different from a random split. Final model evaluation must use official TDC test splits and evaluation metrics as specified per dataset. All experiments should include multiple random seeds to assess reproducibility, and results should be compared against TDC leaderboard baselines available from the TDC website https://tdcommons.ai.

After conducting this research, write a research paper with introduction, results, discussion, methods, and references.

#### Smaller-Scale Prompt

Problem: Evaluating AI-Driven Machine Learning Model Development on Therapeutic Data Commons Benchmarks

The Therapeutic Data Commons (TDC, https://tdcommons.ai) provides standardized benchmarks for drug discovery ML. Run systematic ML experiments to evaluate: (1) what performance levels AI-developed models achieve on TDC datasets compared to leaderboard entries, and (2) what patterns emerge in model failures.

Focus on these 6 TDC datasets: Solubility_AqSolDB, HydrationFreeEnergy_FreeSolv, PAMPA_NCATS, Lipophilicity_AstraZeneca, Bioavailability_Ma, and BBB_Martins. Use the PyTDC ADMET benchmark group API with scaffold splits. For each dataset, find the best ML model and featurization. Run 3 random seeds and report mean and standard deviation. Whenever possible, use splits different from a random split. Final model evaluation must use official TDC test splits and evaluation metrics as specified per dataset. Compare results against TDC leaderboard baselines.

Use the following ML models, in order of priority:

1. Chemprop-RDKit: a graph neural network augmented with 200 RDKit molecular descriptors. Install via pip install chemprop. Documentation and examples at https://github.com/chemprop/chemprop. Use --features_generator rdkit_2d_normalized when running Chemprop-RDKit.
2. Chemprop: the base graph neural network without RDKit features, using the same installation and documentation as above.
3. Other advanced ML models and featurizations, most appropriate for this task, if the above-listed models fail.

After conducting this research, write a research paper with introduction, results, discussion, methods, and references.

### Agent-Based Modeling

#### Initial Prompt

Problem: Improving Price-Setting Behavior in Large-Scale Macroeconomic Agent-Based Models

Agent-based macroeconomic models (ABMs) simulate economies with heterogeneous agents (firms, households, banks, government) interacting through markets. A persistent criticism is that firms in these models rely on overly simplistic, hand-coded heuristics for decision-making, particularly for pricing.

Run advanced scientific research to answer the following research questions:

1. If firms are allowed to learn their pricing strategies through experience rather than following pre-specified rules, what pricing behaviors emerge? Specifically, when firms observe their price relative to sectoral competitors and adjust prices to maximize relative profitability, what strategies do they converge to?
2. Does replacing heuristic pricing rules with learned pricing strategies improve the model’s ability to forecast macroeconomic variables (GDP, inflation, investments, household consumption, government consumption) compared to simple econometric benchmarks?
3. How does the proportion of learning firms affect forecasting performance? Is there an optimal fraction?
4. How do market structure factors (such as whether firm size influences matching probability) affect the learned pricing strategies and their heterogeneity across firms of different sizes and market positions?

Begin by replicating the large-scale, data-driven ABM developed by Poledna et al. (2023, European Economic Review) and validating your replicated model against the calibration results on the Italian economy by Di Domenico et al. (2025, Economic Modelling). This model features heterogeneous firms across multiple industrial sectors, households, a banking sector, and government, with market interactions governed by search-and-matching mechanisms. Use publicly available macroeconomic and sectoral data for the Italian economy, including quarterly national accounts data, Input-Output tables for inter-sectoral linkages, sectoral classification at approximately 60+ industries, firm counts and employment by sector, monetary policy data for the Eurozone. Data should cover a period sufficient for model training, validation, and out-of-sample forecasting. Simulations should be run with thousands of economic agents representing a scaled version of the full economy. Firms should learn independently without centralized coordination. The reward signal should reflect competitive profit maximization (profit relative to sectoral peers).

After conducting this research, write a research paper with introduction, results, discussion, methods, and references.

#### Smaller-Scale Prompt

Study macroeconomic dynamics using an agent-based model of the Italian economy and evaluate how well the model reproduces real economic data. Use the model framework introduced by Poledna et al. (2023, European Economic Review) and the Italian calibration described by Di Domenico et al. (2025, Economic Modelling). Label each modeling decision as empirical, theoretical, or pragmatic and never present assumptions as empirical facts. If multiple modeling choices are possible, explicitly state the uncertainty and explain the chosen option. Use publicly available macroeconomic and sectoral data for the Italian economy, including quarterly national accounts data, Input-Output tables for inter-sectoral linkages, sectoral classification at approximately 60+ industries, firm counts and employment by sector, monetary policy data for the Eurozone. Train on data from 2011-2013, validate on 2014-2016, and test on 2017-2019. Run agent-based simulations representing the Italian economy. Maintain a configuration file that records all model parameters and simulation settings. Record every experiment together with the configuration and random seed so the results can be reproduced. Provide runnable code, configuration files, and scripts for analysis. Produce figures comparing simulated and real macroeconomic time series. Write a research report describing the model, data sources, simulations, and conclusions.

The research should answer the following questions:

How well do the simulated macroeconomic time series such as gross domestic product, inflation, investment, household consumption and government consumption reproduce the empirical dynamics of the Italian economy?

How accurately can the model reproduce or forecast these macroeconomic variables compared with simple statistical benchmarks?

How robust are the simulation results to reasonable variations in parameters, initialization choices and data sources?

